# Evolution Under Thermal Stress Affects *Escherichia coli*’s Resistance to Antibiotics

**DOI:** 10.1101/2024.02.27.582334

**Authors:** Austin Bullivant, Natalie Lozano-Huntelman, Kevin Tabibian, Vivien Leung, Dylan Armstrong, Henry Dudley, Van M. Savage, Alejandra Rodríguez-Verdugo, Pamela J Yeh

## Abstract

Exposure to both antibiotics and temperature changes can induce similar physiological responses in bacteria. Thus, changes in growth temperature may affect antibiotic resistance. Previous studies have found that evolution under antibiotic stress causes shifts in the optimal growth temperature of bacteria. However, little is known about how evolution under thermal stress affects antibiotic resistance. We examined 100+ heat-evolved strains of *Escherichia coli* that evolved under thermal stress. We asked whether evolution under thermal stress affects optimal growth temperature, if there are any correlations between evolving in high temperatures and antibiotic resistance, and if these strains’ antibiotic efficacy changes depending on the local environment’s temperature. We found that: (1) surprisingly, most of the heat-evolved strains displayed a decrease in optimal growth temperature and overall growth relative to the ancestor strain, (2) there were complex patterns of changes in antibiotic resistance when comparing the heat-evolved strains to the ancestor strain, and (3) there were few significant correlations among changes in antibiotic resistance, optimal growth temperature, and overall growth.

**Importance:** *Escherichia coli*, a bacteria species often found within the intestinal tract of warm-blooded organisms, can be harmful to humans. Like all species of bacteria, *E. coli* can evolve, particularly in the presence of stressful conditions such as extreme temperatures or antibiotic treatments. Recent evidence suggests that when encountering one source of stress, an organism’s ability to deal with a different source of stress is also affected. With global climate change and the continued evolution of antibiotic-resistant bacteria, the need to further investigate how temperature and antibiotics interact is clear. The significance of our research is in identifying possible correlations between temperature and antibiotic stress, broadening our understanding of how stressors affect organisms, and allowing for insights into possible future evolutionary pathways.

## Introduction

Environmental stressors such as antibiotics and extreme temperatures can impact the survival and growth of organisms, altering the selection pressures in the environment (Huey & Kingsolver, 1989; Levy & Marshall, 2004; Savage et al., 2004; Bennett & Lenski, 2007; Reed et al., 2011; Buckley & Huey, 2016). In bacteria, these selective pressures can drive the evolution of populations and influence the capacity to withstand perturbations in their environment (Imhof & Schlötterer, 2001; Tello et al., 2012; Van Boeckel et al., 2015; Donhauser et al., 2020). With recent decades seeing a rise in both access to life-saving antibiotics (O’neill, 2014; Van Boeckel et al., 2017; Klein et al., 2018) as well as shifting global temperature due to climate change (Parmesan, 2006; Cavicchioli et al., 2019), understanding the relationships between temperature change and antibiotic exposure is becoming more critical to predicting future trajectories of bacterial populations.

Since life first emerged, organisms have had to evolve mechanisms to survive changes in temperature that stress their physiological processes (Lindquist, 1986; Bada & Lazcano, 2002; Dell et al., 2011; Rohr et al., 2018; Cavicchioli et al., 2019). Both hot and cold temperatures induce different physiological responses in microbes, which impacts their survival and propagation (Yamanaka, 1999; Yura, 2019; Archibald et al., 2022). High temperatures typically result in the misfolding of cellular proteins as well as the formation of aggregates that interfere with essential functions (Richter et al., 2010; Vabulas et al., 2010). To withstand and cope with high-temperature stress, cells have evolved a heat shock response (Ritossa, 1962; Schlesinger et al., 1982), leading to increased expression of chaperone proteins to prevent misfolding as well as proteases to degrade aggregates (Arsène et al., 2000; Roncarati & Scarlato, 2017). Cold temperature responses, on the other hand, are not as thoroughly understood (Yamanaka, 1999). One common response is the stiffening of DNA and RNA structures which results in an overall slower rate of DNA replication and protein synthesis (Phadtare & Inouye, 2008). This stiffening response can also impact lipids, decreasing the efficiency of transport proteins as well as affinity for substrates associated with growth (Yamanaka, 1999; Phadtare & Inouye, 2008; Barria et al., 2013).

Antibiotic compounds have existed long before humans discovered these compounds, however human activities since the 20^th^ century have intensified the strength of their selective pressure (Levy & Marshall, 2004; Mlot, 2009; Davies & Davies, 2010). The more recent widespread use of antibiotics through human activity can also alter bacterial growth and has forced them to adapt to survive in these stressed conditions (Wright, 2005; Souli et al., 2008; Toprak et al., 2012; Van Boeckel et al., 2017; Rodríguez-Verdugo et al., 2020). When evolving resistance to antibiotics, bacteria use one of three main mechanisms to assist in their survival and propagation: (a) *tolerance* allows bacteria to self-inhibit growth when exposed to antibiotics (Kester & Fortune, 2014); (b) *persistence* occurs when a portion of the bacteria population is able to slow growth rates in high drug concentrations (Balaban et al., 2004; Wakamoto et al., 2013); (c) *resistance* is the accumulation of heritable changes that allows bacteria to survive for extended durations in environments with antibiotics (Brauner et al., 2016; Rodríguez-Verdugo et al., 2020). By using these mechanisms, bacteria can often effectively respond to drug-induced stress when faced with various antibiotics.

When an adaptation to one source of stress evolves, it can also alter how a population reacts to other stressors (Brooks & Crowe, 2019; Cruz-Loya et al., 2021). With antibiotics and temperature having some overlap in what cellular functions they affect (Cruz-Loya et al., 2019), it has been suggested that the mechanisms of action between different types of antibiotics and varying temperature ranges may result in cross resistance between the two environmental stressors (Ogbunugafor et al., 2016). Aminoglycosides, for instance, irreversibly associate to the ribosome, and introduce errors in protein translation causing aggregates to form (Mingeot-Leclercq et al., 1999; Ramirez & Tolmasky, 2010; Goltermann et al., 2013). The mechanisms of action for these antibiotics impact similar cellular processes as high temperature conditions (Richter et al., 2010; Vabulas et al., 2010), resulting in non-functional proteins. Further investigation evaluated potential interactions between antibiotics and temperature stress for *E. coli*. Results suggested that aminoglycosides interact with high temperature (46°C) environments to produce higher efficacy at inhibiting bacterial growth than expected from the antibiotics and temperature working alone (Cruz-Loya et al., 2019).

In Tenaillon et al. (2012), derivative strains of an *E. coli* ancestor were evolved under thermal stress for 2,000 generations. While these strains had a wide variety of mutations, adaptive responses to heat stress often resulted specifically in *rpoB* or *rho* termination factor mutations. Some of these strains exhibited a resistance to rifampicin without any prior exposure to antibiotics (Rodríguez-Verdugo et al., 2013). The resistance mutation conferred a considerable selective advantage in stressed environments, and the fitness of the *E. coli* strains varied depending on their genetic background. Results suggest that these bacteria may be employing similar mechanisms to cope with both stressors (Cruz-Loya et al., 2019).

Here we evaluate heat-evolved strains of *E. coli* to determine if there is a correlation between shifts in optimal growth temperature and changes in antibiotic resistance. Specifically, we ask the following questions when *E. coli* is evolved under high temperatures: (1) Do we see any changes in *E. coli*’s optimal growth temperature? (2) Do we see any changes in antibiotic resistance, and, if so, (3) Are these changes correlated?

## Materials and Methods

### Bacterial Strains

We examined the ancestor strain of *Escherichia coli* B genotype *REL1206,* a descendant of *REL606* (Bennett et al., 1992). From a control temperature of 37°C, a total of 114 replicate populations were descended from *REL1206* and independently evolved to a heat stress (42.2°C) for 2,000 generations (Tenaillon et al., 2012).

### Generation of Thermal Performance Curves

We extracted bacteria strains from frozen glycerol stock and allowed them to grow 16 to 18 hours overnight at 37°C. Next, we obtained optical density at 600nm (OD_600_) from each of the overnight samples. We then diluted any strains that displayed an OD_600_ reading above 0.5 by 1:100 in a culture tube with 3mL of fresh lysogeny broth (LB). We allowed any strains below a 0.45 OD_600_ reading to continue growing. After ensuring OD_600_ measurements were consistent across all strains, 200μL of bacteria were transferred into each well of a 96-well plate. We divided the 96-well plate in half to provide six replicates of thirteen strains for more accurate results. The bacteria were then pin-transferred into 96-well plates containing 200μL of LB per well before being covered by a porous seal to allow for gas exchange. We placed the plates in incubators with sixteen staggered temperatures ranging from 12°C to 50°C for approximately 18 to 22 hours. OD_600_ measurements were collected after the incubation period. We used six replicates of LB only as a negative control and subtracted from the OD_600_ of the sample strains to only measure bacterial growth.

Our goal was to understand how each strain grew over a wide range of temperatures. To generate thermal performance curves for each strain, we used the methods described by Cruz-Loya et al. (2021). In summary, we used a modified Briere function and defined the temperature dependence of growth as g(T). The reparametrized extended Briere model is as follows:

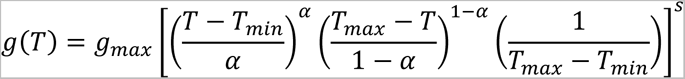

where the value of *g_max_* is the highest measured growth—the highest OD_600_ value after approximately 20hr—of the examined strain, and values of *α* and s alter the shape of the bacterial growth curve (Cruz-Loya et al., 2021). In this reparametrized equation, the first term within the brackets enforces growth to go to zero at a minimum temperature, the second term enforces growth to go to zero at a maximum temperature, and the third term along with the values of *α* and *s* accounts for the full range of growth and the location of the optimal temperature for a given strain of *E. coli*. We then calculated the optimal growth temperature (T_opt_) for each strain using:

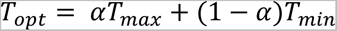

We used the function nlsLM within the minpack.lm package version 1.2-4 in R to fit our models (Elzhov et al., 2016). Due to the nature of the reparametrized Briere model, data with negative or zero growth values were removed from our data set during model fitting. A few examples of fitting this model to our data set can be found in Figure 1. This model was unable to fit the growth data for six strains (Strains 32, 36, 85, 108, 118, and 119). These strains were then removed from the analysis.

**Figure 1.**
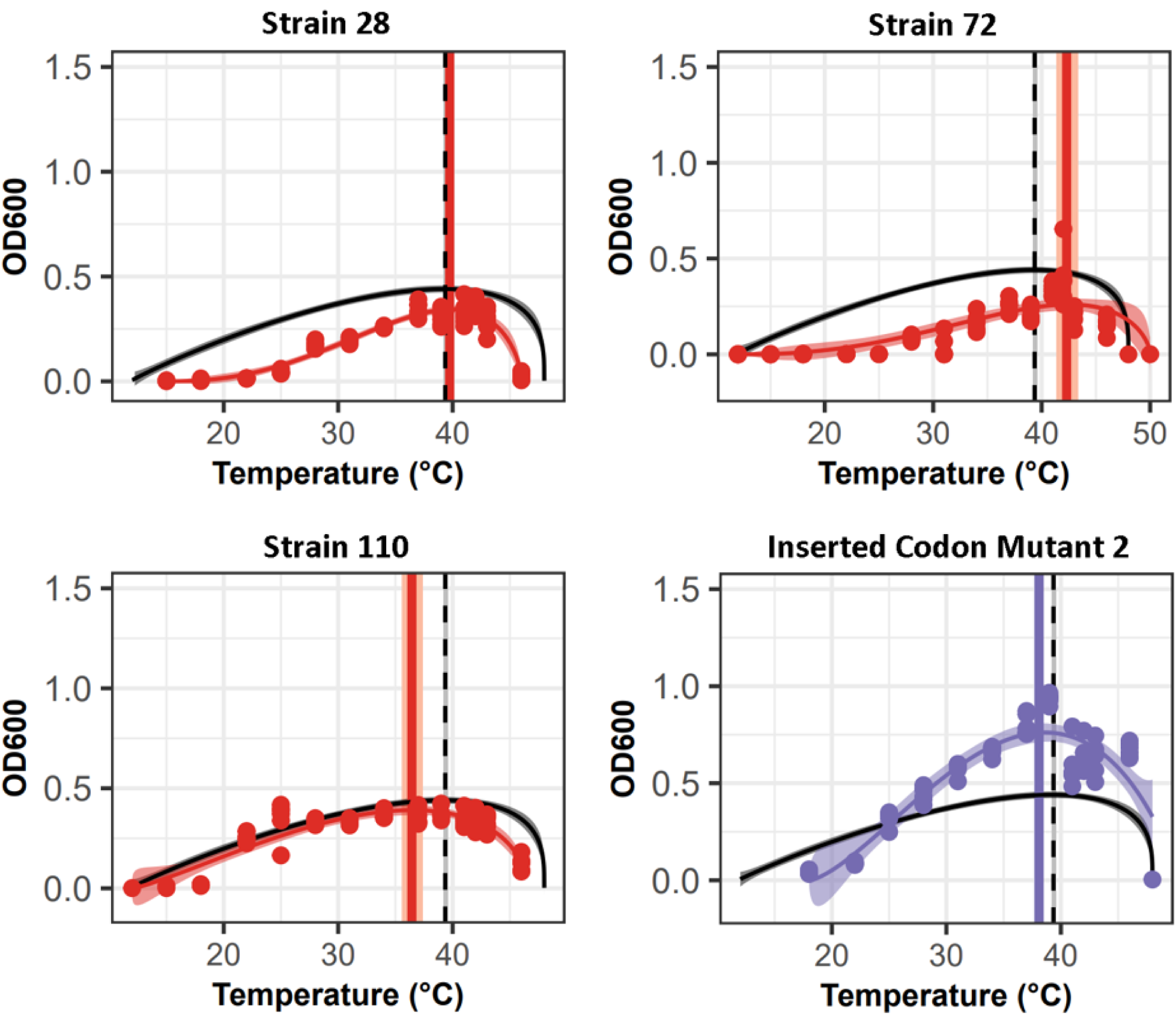
Examples of the thermal performance curves (TPC) and calculated optimal temperatures fitted to data with the reparametrized Briere model from Cruz-Loya et al. (2021). The solid black line shows the TPC of the ancestral strain of *E. coli* with the vertical, dashed black line at the ancestral optimal growth temperature. Solid red or purple lines show the heat-evolved (red) or mutant (purple) strain’s TPC while the respective vertical line show the optimal temperature. 95% confidence intervals for both the model and optimal temperature are displayed as shaded regions. Strains were selected to be representatives of the optimal growth temperature shifts observed.

Once we generated heat response curves with the modified Briere model, we calculated the area under the curve between T_min_ and T_max_ for each strain using the AUC function within the DescTools package version 0.99.5 in R (Signorell et al., 2019). Additionally, we calculated the difference in area between the ancestral strain’s heat response curve and each individual heat-evolved strain’s response curve to evaluate overall growth.

### Determining Strength of Antibiotic Resistance

To test for antibiotic resistance, we ran a series of serial dilutions for each antibiotic in 96-well plates. Specifically, we diluted the antibiotics twenty times with each dilution being one half of the previous concentration. We used twelve antibiotics for our experiments that span the major classifications of antibiotics used in medical settings (Table 1). We extracted antibiotics from a stock solution and diluted in LB to a concentration of 4000μg/mL. We added antibiotics into a 96-well plate containing 100μL of LB and serially diluted wells in the plate, establishing a range of concentrations from 2000μg/mL to approximately 0.008μg/mL. We adjusted the starting concentration values for antibiotics that required greater resolution to determine antibiotic resistance. We sealed the plates and placed them in a 37°C incubator for 22–24 hours before we collected OD_600_ measurements. We determined and compared IC_50_ values, the minimum amount of antibiotic needed to kill 50% of the bacteria population, to evaluate antibiotic resistance. The IC_50_s for all heat-evolved strains were determined when plates were incubated at 37°C, corresponding to a non-stressed environment. We also took a small subsample of fifteen random strains to determine the IC_50_ data for each antibiotic at 42°C—the evolved environment—to assess how this may change antibiotic resistance.

**Table 1.**
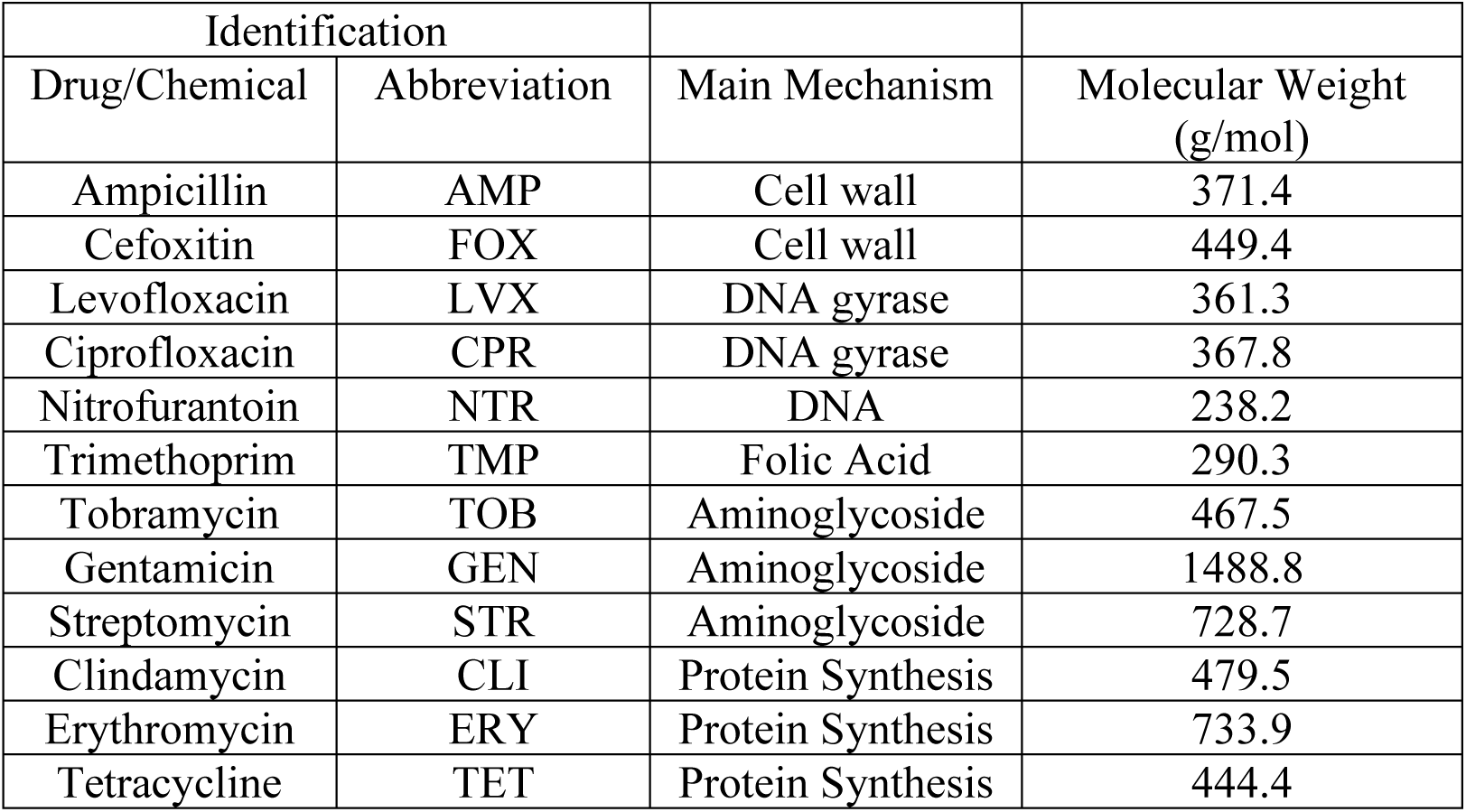
List of antibiotics examined in this study.

When testing the heat-evolved strains against antibiotics, some strains grew beyond 2000 μg/mL. Those samples had their IC_50_ values set to 2000 μg/mL to minimize errors generating MIC figures.

We determined the log_2_ mean fold change of the IC_50_ to evaluate how evolving in high temperatures potentially changed the IC_50_ value and thus antibiotic resistance levels.

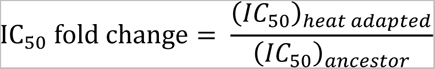

Values greater than one indicate that heat adaptation also conferred resistance to the antibiotic and values less than one indicate heat adaptation increased sensitivities to the antibiotic.

## Results

### Heat-Evolved Strains Display Decreased Optimal Growth Temperature and Decreased Overall Growth

We found optimal growth temperature tended to decrease for the heat-evolved strains with an overall average of 36.2°C (std. dev. = 2.4°C). This is a significant decrease compared to the optimal growth temperature of the ancestral strain (37.92°C) (two-tailed, one-sample t-test, *μ* = 37.92, p = 5.9 × 10^−11^). However, 23 strains out of 108 strains did have a higher optimal growth temperature relative to the ancestor (Figure 2A). We observed a wide variety of optimal growth temperatures, with some showing relatively little change (Figure 1 Strain 28), few showing an increase (Figure 1 Strain 72), and most showing a decrease (Figure 1 Strain 110). The optimal temperature is only one point of comparison. To compare the overall shape of the temperature responses more holistically across the full range, we measured the difference in area between heat response curves of the ancestral and heat-evolved strains. We observed that a majority of our heat-evolved strains (∼96%) displayed lower overall growth area across the tested temperature range (Figure 2B). Examples of this decreased overall growth can be found in Figure 1 (Strain 28, 72, and 110). Additionally, we found that comparing changes in optimal growth temperature with changes in the area between heat response curves yielded a significant negative relationship between the two variables, suggesting that these are not independent effects (Spearman correlation, R= 0.37, p < 0.0001) (Figure 3).

**Figure 2.**
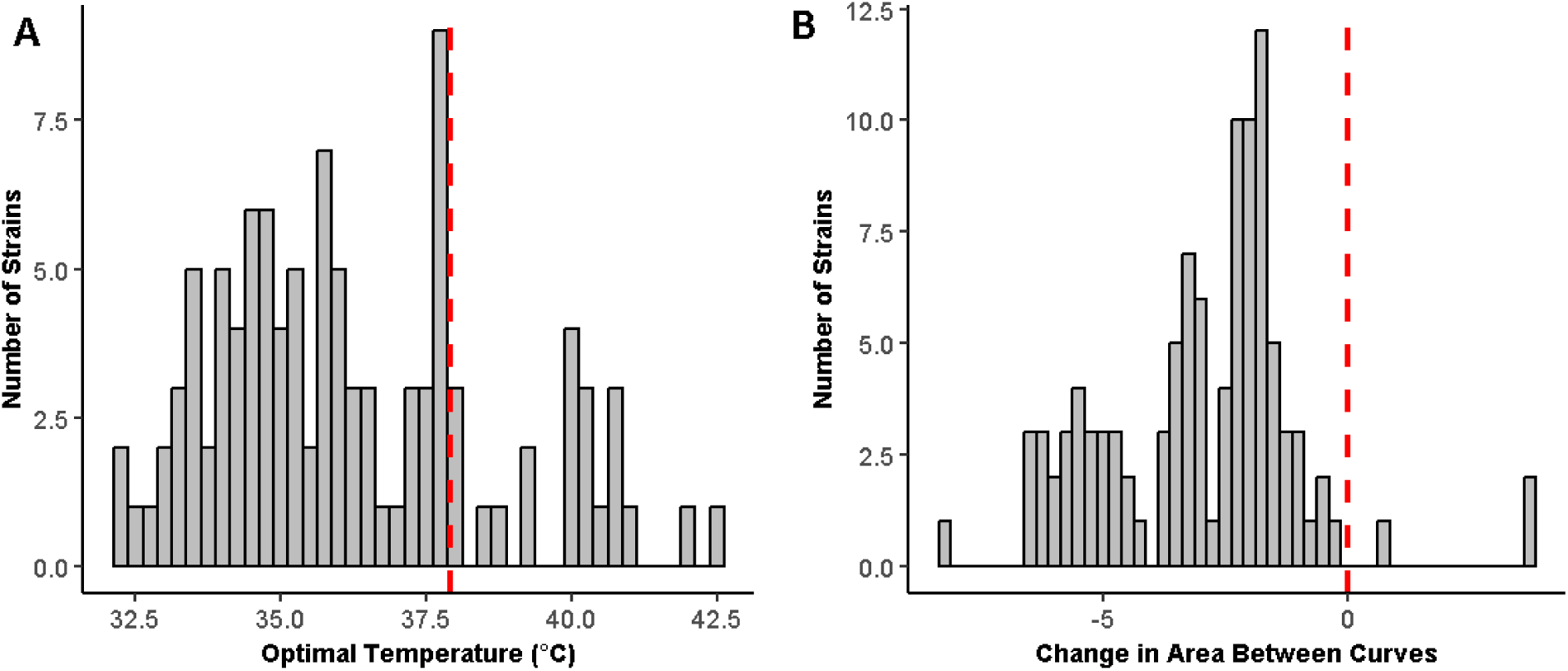
Distribution of optimal temperature and change in overall growth for all strains examined. (A) The majority of the heat-evolved strains (∼79%) had a lower optimum temperature compared to the ancestor (37.92°C). The ancestral optimal growth temperature is shown as a red dashed line. (B) Nearly all (96%) of the examined strains had less overall growth when evaluating the area under the curve of the thermal performance curves generated. The vertical, red dashed line represents area under the ancestral thermal performance curve. The change in area between curves is the difference between a heat-evolved strain and the ancestral strain. Negative values reflect an instance where the ancestral strain demonstrated higher amounts of bacterial growth across the tested temperatures, whereas positive values were the result of a heat-evolved strain showing higher growth than the ancestral strain.

**Figure 3.**
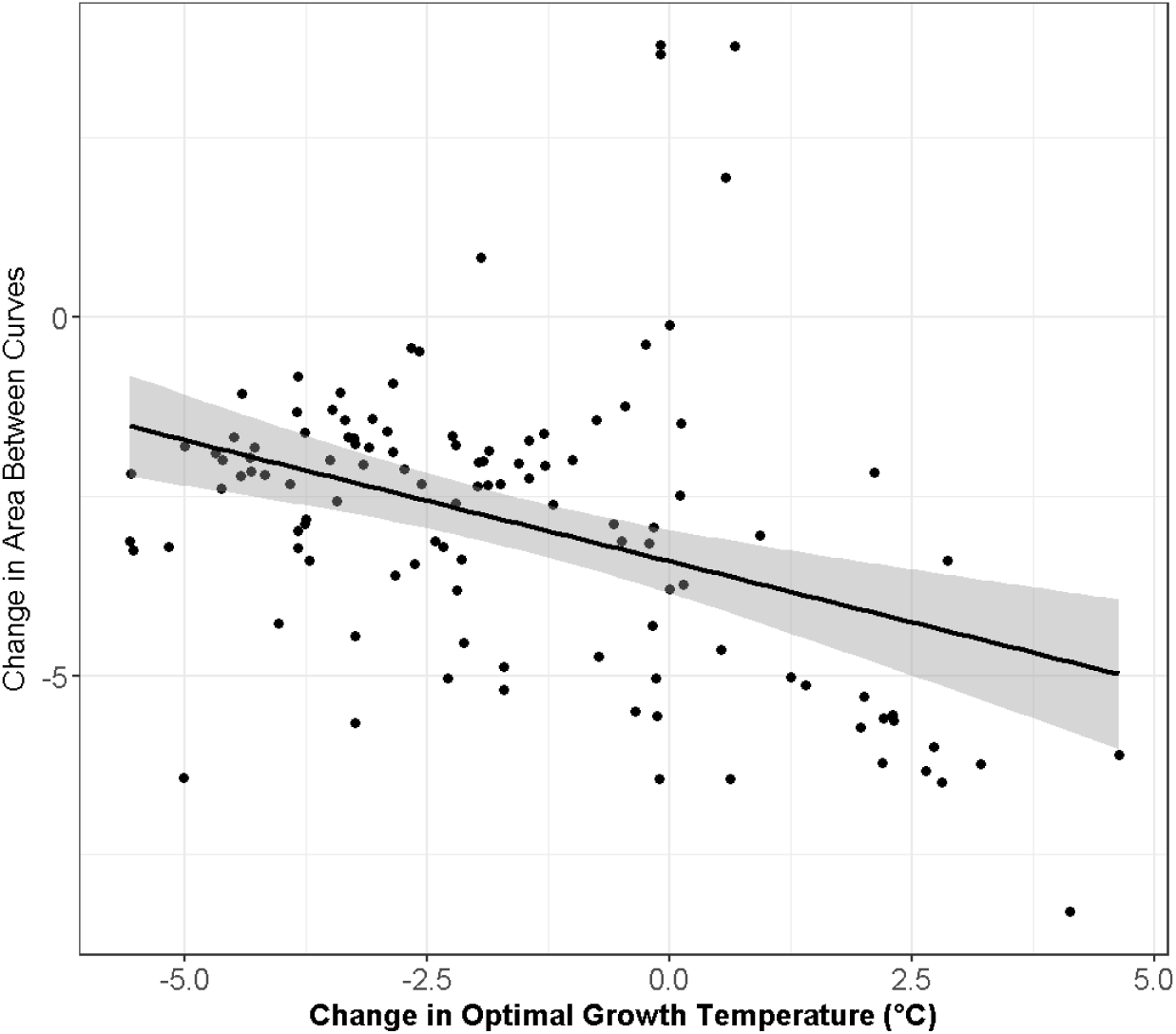
A negative correlation between the area between heat response curves and change in optimal growth temperature of the heat-evolved strains (Spearman correlation, R = −0.37, p = 5.9 × 10^−5^).

### Strength of Antibiotic Resistance Changed between Different Temperatures

In general, evolving in high temperatures changed the strains’ IC_50_ values when exposed to several different antibiotics (Figure 4). We found a significant increase of resistance to gentamicin and levofloxacin when IC_50_ values were determined at 37°C. However, only levofloxacin showed a significant increase in IC_50_ value at both 37°C and 42°C (two-tailed, one-sample t-test, *μ* = 1, p_37°C_ = 9.47 × 10^−4^, p_42°C_ = 1.14 × 10^−7^) (SI Table 1 and SI Table 2). We also found a significant increase in sensitivity to ampicillin, ciprofloxacin, cefoxitin, clindamycin, tetracycline, and trimethoprim (two-tailed, one-sample t-test, *μ* = 1, p < 2.73 × 10^−4^, SI Table 1) when IC_50_ values were determined at 37°C. This significant increase in sensitivity was also found in ampicillin, clindamycin, ciprofloxacin, and tetracycline when IC_50_ values were determined at 42°C (two-tailed, one-sample t-test, *μ* = 1, p < 1.54 × 10^−7^, SI Table 2).

**Figure 4.**
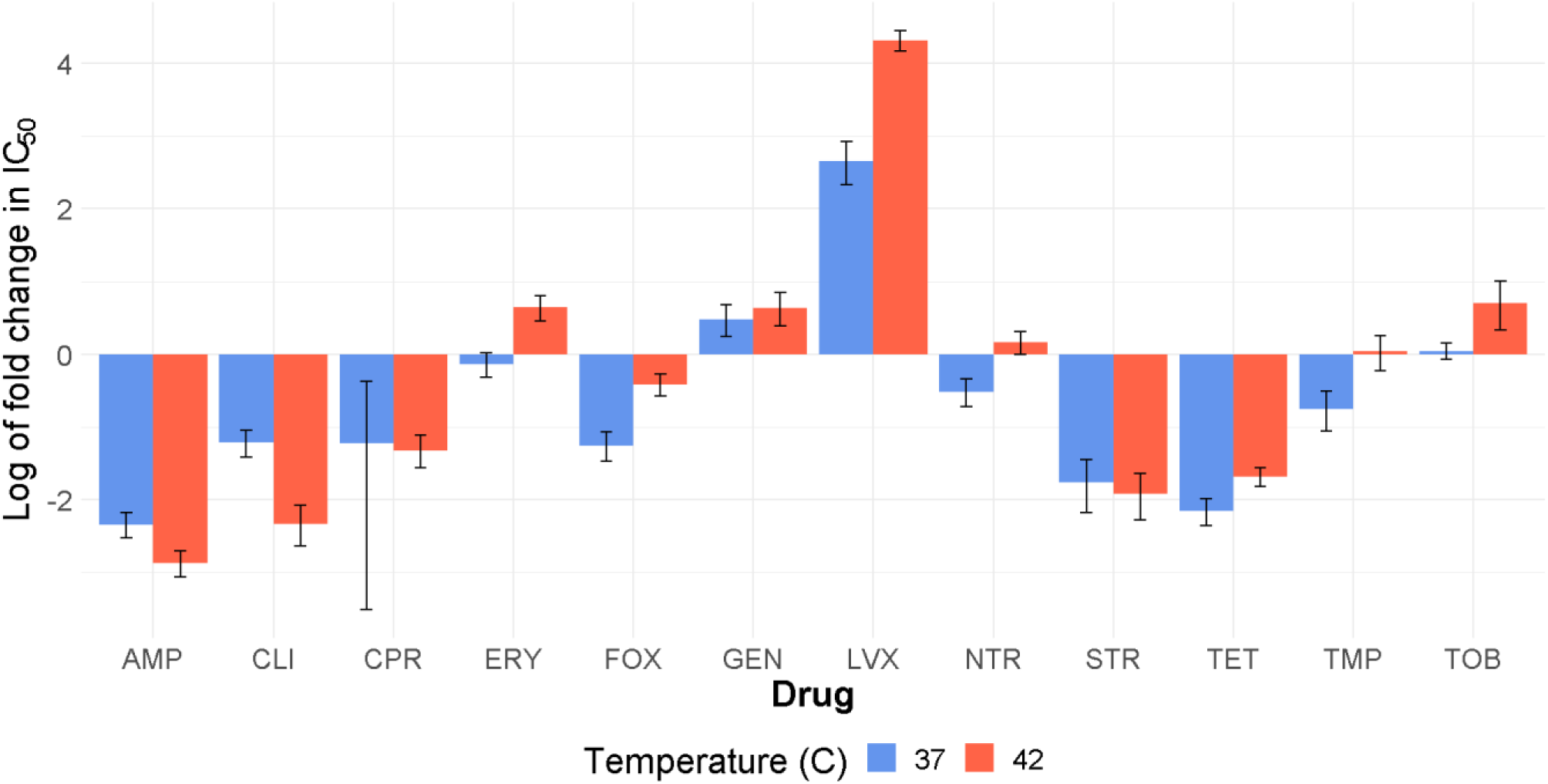
The fold change of antibiotic resistance 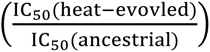 of the *E. coli* strains at 37°C (blue) and 42°C (red), typically varied or lower resistance. Log_2_ of fold change was used to compare the heat-evolved strains between the two temperature conditions with the ancestor strain’s antibiotic resistance. Error bars show standard mean error. The dose response curve for individual strains for ciprofloxacin (CPR) displayed a large variation in IC_50_ values.

**Table 2.** The correlation strength (R) and p-value of Spearman correlation between IC_50_ values and optimal growth temperature of the heat-evolved strains. Significant correlations (p < 0.05) are shown in **bold**. These results show that IC_50_ values and optimal growth temperature are mostly independent of one another across antibiotics.

| Drug | Temperature (°C) | # of Strains | Spearman Correlation (R) | p-value |
| --- | --- | --- | --- | --- |
| AMP | 37 | 108 | -0.061 | 0.55 |
| CLI | 37 | 108 | 0.12 | 0.23 |
| CPR | 37 | 108 | 0.025 | 0.81 |
| **ERY** | **37** | **108** | **0.2** | **0.047** |
| FOX | 37 | 108 | 0.052 | 0.61 |
| GEN | 37 | 108 | -0.0035 | 0.73 |
| LVX | 37 | 108 | -0.13 | 0.19 |
| **NTR** | **37** | **108** | **0.22** | **0.032** |
| STR | 37 | 108 | 0.099 | 0.45 |
| TET | 37 | 108 | -0.0042 | 0.97 |
| TMP | 37 | 108 | 0.17 | 0.094 |
| TOB | 37 | 108 | 0.12 | 0.23 |
| AMP | 42 | 15 | -0.14 | 0.63 |
| CLI | 42 | 15 | 0.17 | 0.54 |
| CPR | 42 | 15 | 0.14 | 0.62 |
| ERY | 42 | 15 | 0.11 | 0.7 |
| FOX | 42 | 15 | 0.086 | 0.76 |
| **GEN** | **42** | **15** | **0.61** | **0.017** |
| LVX | 42 | 15 | 0.42 | 0.12 |
| NTR | 42 | 15 | 0.34 | 0.21 |
| STR | 42 | 15 | 0.16 | 0.58 |
| TET | 42 | 15 | -0.027 | 0.93 |
| TMP | 42 | 15 | 0.14 | 0.63 |
| **TOB** | **42** | **15** | **0.66** | **0.0086** |

We compared the IC_50_ of each antibiotic at 37°C and 42°C to see if there were any differences in antibiotic resistance between the heat-evolved strains and the ancestral strain (Figure 4). We found that three of the twelve antibiotics (cefoxitin, clindamycin, and levofloxacin) showed significantly different IC_50_ values at 42°C compared to 37°C (SI Table 3). Levofloxacin and cefoxitin showed significantly higher IC_50_ values at 42°C (two-tailed, two-sample t-test, p < 0.003). IC_50_ values for clindamycin was significantly higher at 37°C (two-tailed, two-sample t-test, p= 0.002). A full listing of p-values and test statistics can be found in SI Table 3.

**Table 3.** The correlation strength (R) and p-value of Spearman correlation between IC_50_ values and changes in area for heat response curves for the heat-evolved strains. Significant correlations (p < 0.05) are shown in **bold**. Only nitrofurantoin (NTR) exhibits a significant correlation between IC_50_ values and changes in area between heat response curves. This indicates that these variables are mostly independent.

| Drug | Temperature (°C) | # of Strains | Spearman Correlation (R) | p-value |
| --- | --- | --- | --- | --- |
| AMP | 37 | 108 | 0.0056 | 0.96 |
| CLI | 37 | 108 | 0.078 | 0.44 |
| CPR | 37 | 108 | 0.0013 | 0.99 |
| ERY | 37 | 108 | -0.1 | 0.31 |
| FOX | 37 | 108 | -0.069 | 0.5 |
| GEN | 37 | 108 | -0.037 | 0.72 |
| LVX | 37 | 108 | -0.082 | 0.42 |
| **NTR** | **37** | **108** | **-0.25** | **0.013** |
| STR | 37 | 108 | 0.19 | 0.14 |
| TET | 37 | 108 | 0.11 | 0.29 |
| TMP | 37 | 108 | -0.014 | 0.89 |
| TOB | 37 | 108 | -0.083 | 0.42 |
| AMP | 42 | 15 | 0.21 | 0.46 |
| CLI | 42 | 15 | 0.032 | 0.91 |
| CPR | 42 | 15 | 0.2 | 0.47 |
| ERY | 42 | 15 | -0.089 | 0.75 |
| FOX | 42 | 15 | 0.45 | 0.094 |
| GEN | 42 | 15 | -0.23 | 0.4 |
| LVX | 42 | 15 | -0.064 | 0.82 |
| NTR | 42 | 15 | 0.029 | 0.92 |
| STR | 42 | 15 | 0.16 | 0.59 |
| TET | 42 | 15 | 0.016 | 0.96 |
| TMP | 42 | 15 | 0.39 | 0.16 |
| TOB | 42 | 15 | -0.27 | 0.32 |

### Few Significant Relationships between Optimal Growth Temperature, Overall Growth, and Antibiotic Resistance

We investigated whether there are any relationships between changes in optimal growth temperature and corresponding IC_50_ values. We observed a significant correlation between IC_50_ values and optimal growth temperature changes for erythromycin at 37°C, nitrofurantoin at 37°C, gentamicin at 42 °C, and tobramycin at 42°C (Spearman correlation, p < 0.05, Table 2). The correlation between the tobramycin IC_50_ values determined at 42°C and the change in optimal growth temperature showed the strongest correlation, which was a positive correlation (as optimal temperature increased so did the IC_50_ value for tobramycin) (Spearman correlation, R= 0.66, p= 0.0086) (Figure 5A).

**Figure 5.**
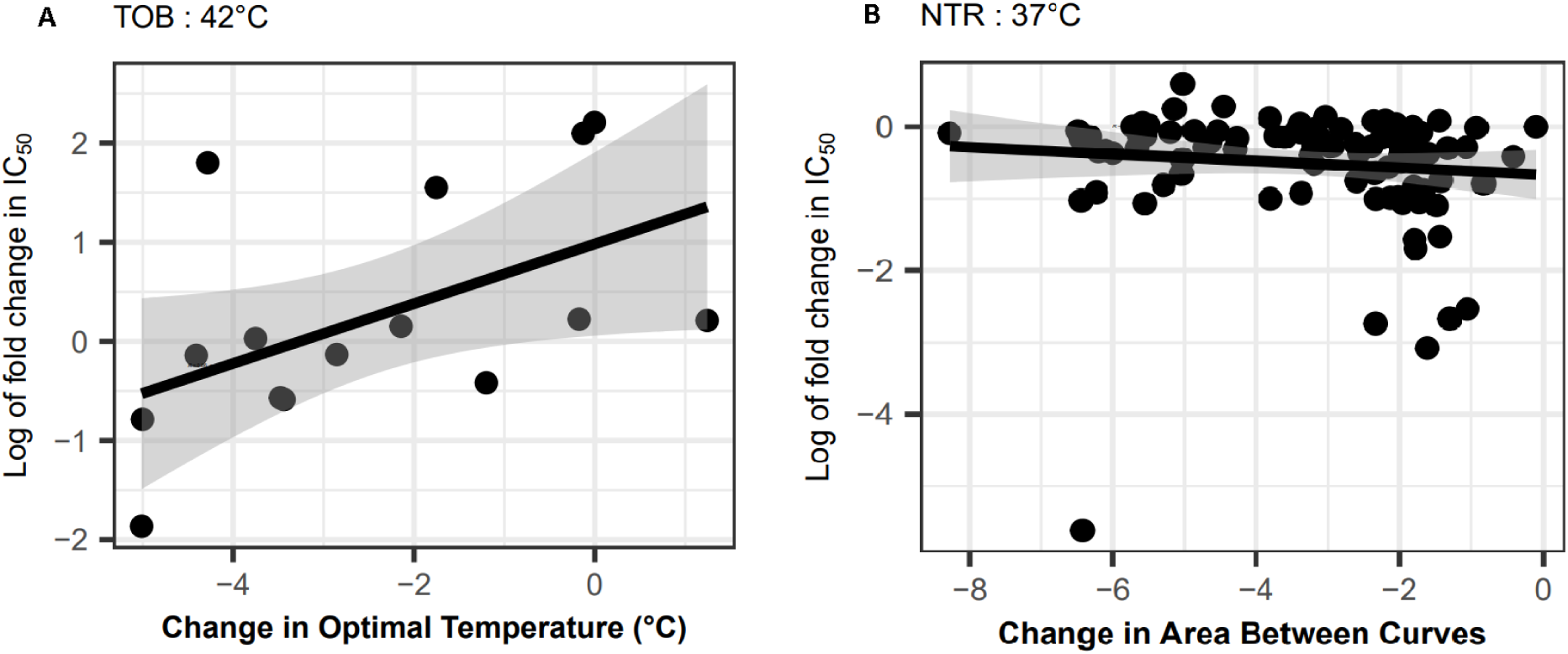
The two most significant correlations between antibiotic resistance and change in optimal temperature or overall growth. (A) A representative of antibiotic resistance evaluated at 42°C when correlated with changes in optimal temperature. We found a positive correlation between tobramycin (TOB) resistance and increasing optimal temperature (Spearman correlation, R = 0.66, p = 0.0086, Table 2). (B) Nitrofurantoin (NTR) had the only significant correlation when comparing IC_50_ values and changes in overall growth when examined at 37°C (Spearman correlation, R = −0.25, p = 0.013, Table 3).

Further examining our strains, we observed that only nitrofurantoin at 37°C had a significant correlation between IC_50_ values and changes in area between heat response curves (Spearman correlation, R= −0.25, p= 0.013, Figure 5B Table 3).

## Discussion

We asked how adaptation to higher thermal stress on *E. coli* may affect their optimal growth temperature, overall growth, and antibiotic resistance. We found that optimal growth temperature varied among the 100+ strains, with most showing a lower optimal growth temperature than the ancestor strain’s optimum temperature. No strains exhibited a significant relationship between antibiotic resistance and changes in optimum growth temperature, however we observed a significant negative relationship when comparing the area between heat response curves and optimal growth temperature. We did see significant changes in antibiotic resistance between the heat-evolved strains and the ancestor strain.

We hypothesized that the evolved strains examined would have an increased optimal growth temperature relative to the ancestral strain (Rodríguez-Verdugo et al., 2014). Heat shock responses are a highly conserved physiological response among prokaryotes and eukaryotes (Richter et al., 2010; Hug & Gaut, 2015). This response promotes increased synthesis of heat-shock proteins and chaperones to degrade any formed aggregates and prevent further protein misfolding (Vabulas et al., 2010; Mondal et al., 2014). When exposed to high temperatures for long periods of time, individuals that show increased expression of this heat-shock response will likely be favored in this stressed environment (Deatherage et al., 2017). Experimental evolution of bacteria under thermal stress can adjust in response to stress through two possible strategies. In the short-term, phenotypic plasticity can help bacteria acclimate whereas genetic changes help in longer-term scenarios (Hug & Gaut, 2015). With our heat-evolved strains experiencing high heat stress for 2,000 generations, we expected to see an increased optimal growth temperature (Rodríguez-Verdugo et al., 2014). However, our results did not match our initial hypothesis. We were surprised to find that the majority (79%) of our heat-evolved strains displayed a lower optimal growth temperature than the ancestral strain (Figure 2A).

One possible explanation for the decreased optimal growth temperatures seen in many of the heat-evolved strains could be due to an interaction between adaptation and acclimation to the bacteria’s local environment (Hug & Gaut, 2015). A previous study investigated how the phenotypes of heat-evolved *E. coli* would respond when placed in different temperature conditions. The majority of the heat-evolved strains’ phenotypes displayed a return to the unstressed phenotypic state (Hug & Gaut, 2015). Our observations of lower optimal growth temperatures for *E. coli* support previous studies, which suggest *E. coli* can employ adaptive strategies in an attempt to return the bacterium cell to its unstressed physiological state found at 37°C (Hug & Gaut, 2015; Lambros et al., 2021). An attempt to return to an unstressed physiological state may enable bacteria to increase their fitness in stressful environments (Lambros et al., 2021).

Another possible explanation for the observed decreases in optimal growth temperatures is that these heat-evolved have high phenotypic plasticity for temperature stress (Miller et al., 2020). A previous study focusing on how cyanobacteria respond to extreme temperature argued that plasticity allows for innovation to arise in stressed populations (Levis & Pfennig, 2020; Miller et al., 2020). It was found that under extreme temperature conditions some cyanobacteria populations developed a novel form of a cell wall that is less permeable allowing them to survive in these high temperatures (Miller et al., 2020). It is thought that this plasticity allows the bacteria to “buy time” in novel environments and helps them to persist in conditions that may not be optimal (Fox et al., 2019).

We observed that, in addition to many of the heat-evolved strains (96%) displaying lower growth levels than the ancestral strain across the tested temperature range (Figure 2B), there is a significant negative relationship between changes in optimal growth temperature and changes in area between heat response curves (Figure 3). One plausible explanation for this correlation could be that bacteria that are exposed to a novel temperature environment accumulate mutations that negatively impact their fitness (Bosshard et al., 2017). The temperature ranges that the heat-evolved strains were exposed to likely introduced mutations that, while allowing the strains to grow in more stressful temperature conditions, resulted in weaker bacterial growth overall. Interestingly, two of the three mutant strains that had a direct insertion of a *rpoB* mutation displayed greater bacterial growth compared to the ancestral strain (Rodríguez-Verdugo et al., 2014) (Figure 1, Inserted Codon Mutant 2). This insertion of the mutation may have some role in preventing the deleterious effects of novel mutations acquired by the heat-evolved strains as they were evolved under thermal stress while conferring a fitness benefit to the mutant strains (Bosshard et al., 2017).

We also hypothesized that antibiotic resistance for our heat-evolved strains should change depending on the temperature conditions they are grown under. Cruz-Loya et al. (2019) highlighted a stressor network between temperature and antibiotics. This is in part due to how “hot-like” antibiotics, such as aminoglycosides, can negatively affect the same cellular mechanisms that hot temperatures do (Goltermann et al., 2013; Cruz-Loya et al., 2019). Further observations on possible interactions between antibiotics and stressors can also have potential clinical applications as prior work has found that determining bacterial resistomes can be useful in designing effective codrugs (Liu et al., 2010).

We observed significant increases in antibiotic resistance (IC_50_) for multiple antibiotics. The heat-evolved strains had significantly different levels of resistance to the majority (10 of the 12) of the antibiotics we tested at 37°C (SI Table 1). Among these ten, six of them had significantly different resistances at 42°C (SI Table 2). This suggests that heat-adaptation affected resistance levels to these antibiotics. Interestingly, one of the drugs that our strains were most resistant to at 37°C and 42°C was levofloxacin (LVX) which is associated with greater performance at cold temperatures (Cruz-Loya et al., 2019). Also, despite having similar mechanisms of action, not every antibiotic of the same class responded to temperature in the same manner.

We hypothesized that for heat-evolved *E. coli* strains, there are relationships between changes in optimal growth temperature, overall bacterial growth, and strength of antibiotic resistance. However, our results did not support our initial hypothesis. With some antibiotic classes and temperature overlapping in the cellular mechanisms they affect, an adaptation to temperature may confer increased fitness against antibiotics (Cruz-Loya et al., 2019). Prior studies have shown that, if possible, cells will attempt to evolve a co-opted response that is able to function when exposed to various types of stressors (Dragosits et al. 2013; Święciło 2016). However, in this study, we found few significant correlations between optimal growth temperature, overall growth, and antibiotic resistance (Table 2, Table 3).

One possible explanation as to why we see few significant relationships between optimal growth temperature, overall growth, and antibiotic resistance may be that their novel traits do not necessarily provide an immediate fitness benefit (Karve et al., 2015; Toll-Riera et al., 2016). Novel mutations that are not the primary focus of experimental evolution can arise in bacterial populations over time (Karve & Wagner, 2022). These novel mutations can appear as a byproduct of the evolution of other adaptive traits and may become beneficial once the environment changes (Karve & Wagner, 2022). Fitness trade-offs have been observed when bacteria are evolved under antibiotic stress before exposure to novel temperatures (Herren & Baym, 2022). Strains that evolved resistance to antibiotics were found to have reduced growth at extreme temperature ranges and relatively normal growth at optimal growth temperature conditions (Herren & Baym, 2022). Our results suggest that in unstressed temperature environments, changes in antibiotic resistance might not be directly due to shifts in optimal growth temperature.

Other studies that have focused on the effects of temperature stress have examined temperature niche breadth, focusing on a range of temperatures where bacteria can grow effectively (MacFadden et al., 2018; Herren & Baym, 2022). Examining a range of temperatures may provide useful information for examining heat-adaptation, such as any phenotypic plasticity that bacteria can exhibit in response to temperature stress (Payne & Wagner, 2019). Although we examined a range of temperatures to determine the optimal growth temperature of the heat-evolved *E. coli,* a more thorough examination of niche temperature breadth would provide a more complete understanding of how different dimensions of heat adaptation affect antibiotic resistance.

With increases in temperature forecast in many parts of the world as a result of global climate change, it is becoming more crucial to understand temperature-antibiotic interactions. Despite numerous studies on how climate change can affect disease vectors (Campbell-Lendrum et al., 2015; Ogden & Lindsay, 2016; Castledine et al., 2020), few have examined how the pathogens themselves would be affected by shifts in temperature (Rodríguez-Verdugo et al., 2020). Novel temperature environments can affect bacteria by spatially shifting antibiotic-resistant populations across different geographic regions (MacFadden et al., 2018), adjusting the temperature range at which these organisms can grow (Knies et al., 2009), and also lead to increased rates of antibiotic resistance (Ratkowsky et al., 1982; McGough et al., 2020). Our work shows how exposing *E. coli* to long-term heat stress can yield a range of responses for optimal growth temperature and antibiotic resistance. Many of the replicate *E. coli* strains were found to have lower optimal growth temperatures relative to the ancestral strain and among the tested antibiotics, two-thirds of them yielded significantly different resistance levels between the 37°C and 42°C environments. With an actively changing global climate, understanding potential patterns between stressors may help us better predict evolutionary trajectories of future bacterial populations.

## Data Availability

MIC and heat response data, as well as the code, and figures used in this paper will be available in the Mendeley Data Repository upon publication of the article (Bullivant, 2024).

## Acknowledgements

We thank Emoni Cook and Nicholas Ida for assistance in the laboratory as well as Colin Kremer for suggestions on the manuscript. The research described was supported by NIH/National Center for Advancing Translational Science (NCATS) UCLA CTSI Grant Number UL1TR001881.

## Supplemental Information

**SI Table 1.**
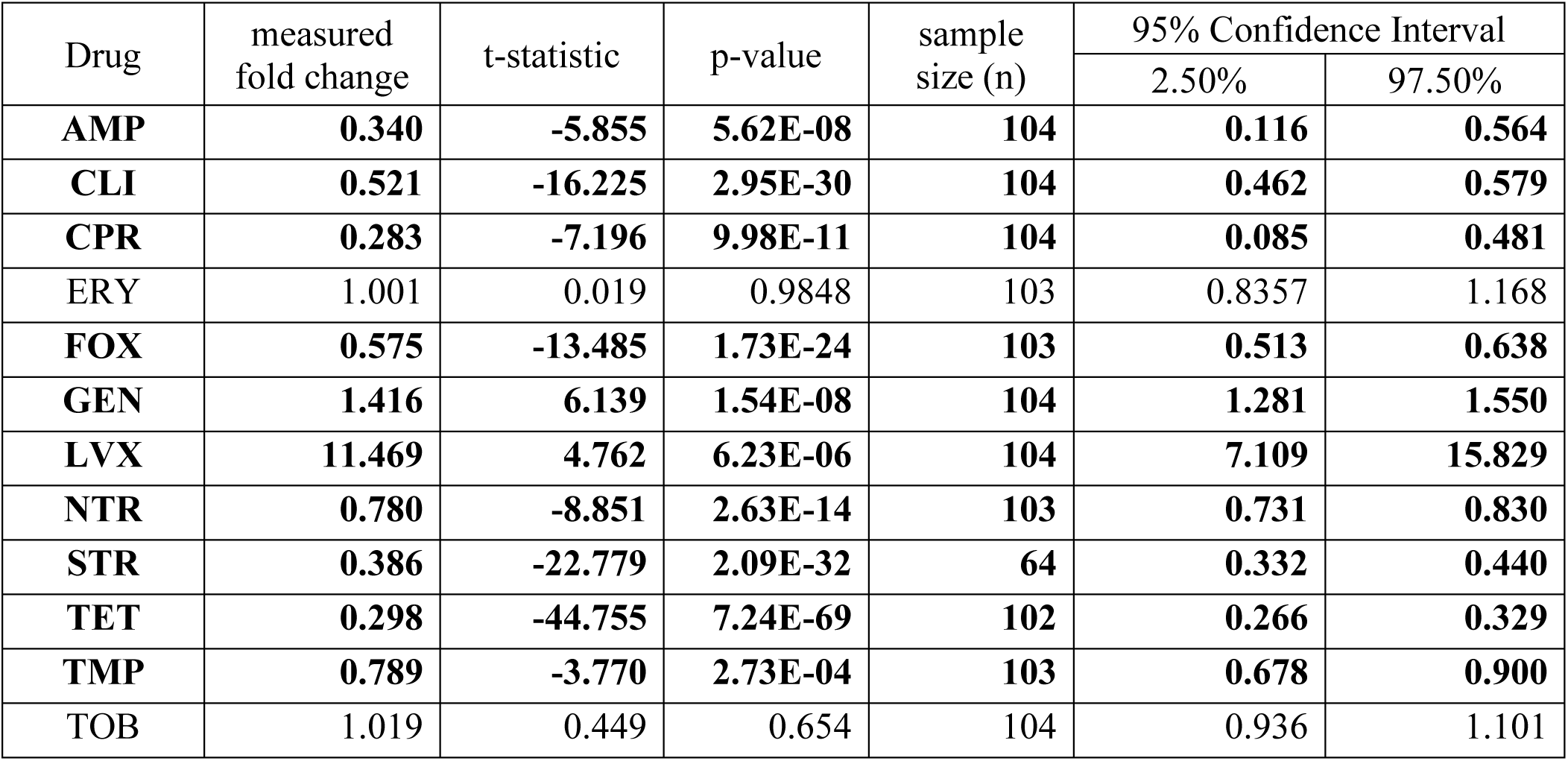
The fold change of IC_50_ values after heat evolution at 42.2℃ 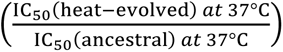 when in a 37°C environment. Comparisons in **bold** indicate a significant difference—meaning the fold change is statistically significantly different from 1 (indicating no change in response as compared with the ancestral strain)—after performing a Bonferroni multiple comparison correction (Bonferroni corrected *α* = 0.002).

| Drug | measured fold change | t-statistic | p-value | sample size (n) | 95% Confidence Interval |  |
| --- | --- | --- | --- | --- | --- | --- |
|  |  |  |  |  | 2.50% | 97.50% |
| **AMP** | **0.340** | **-5.855** | **5.62E-08** | **104** | **0.116** | **0.564** |
| **CLI** | **0.521** | **-16.225** | **2.95E-30** | **104** | **0.462** | **0.579** |
| **CPR** | **0.283** | **-7.196** | **9.98E-11** | **104** | **0.085** | **0.481** |
| ERY | 1.001 | 0.019 | 0.9848 | 103 | 0.8357 | 1.168 |
| **FOX** | **0.575** | **-13.485** | **1.73E-24** | **103** | **0.513** | **0.638** |
| **GEN** | **1.416** | **6.139** | **1.54E-08** | **104** | **1.281** | **1.550** |
| **LVX** | **11.469** | **4.762** | **6.23E-06** | **104** | **7.109** | **15.829** |
| **NTR** | **0.780** | **-8.851** | **2.63E-14** | **103** | **0.731** | **0.830** |
| **STR** | **0.386** | **-22.779** | **2.09E-32** | **64** | **0.332** | **0.440** |
| **TET** | **0.298** | **-44.755** | **7.24E-69** | **102** | **0.266** | **0.329** |
| **TMP** | **0.789** | **-3.770** | **2.73E-04** | **103** | **0.678** | **0.900** |
| TOB | 1.019 | 0.449 | 0.654 | 104 | 0.936 | 1.101 |

**SI Table 2.** The fold change of IC_50_ values after heat evolution at 42.2℃ 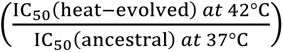 when in a 42°C environment. Comparisons in **bold** indicate a significant difference—meaning the fold change is statistically significantly different from 1 (indicating no change in response as compared with the ancestral strain)—after performing a Bonferroni multiple comparison correction (Bonferroni corrected *α* = 0.002).

| Drug | measurement | t-statistic | p-value | sample size<br>(n) | 95% Confidence Interval |  |
| --- | --- | --- | --- | --- | --- | --- |
|  |  |  |  |  | 2.50% | 97.50% |
| **AMP** | **0.137** | **-51.157** | **2.54E-17** | **14** | **0.101** | **0.173** |
| **CLI** | **0.199** | **-21.199** | **4.88E-12** | **14** | **0.117** | **0.280** |
| **TET** | **0.311** | **-25.497** | **8.06E-12** | **12** | **0.253** | **0.370** |
| **CPR** | **0.400** | **-9.608** | **1.53E-07** | **14** | **0.267** | **0.534** |
| FOX | 0.749 | -3.260 | 0.006 | 14 | 0.583 | 0.914 |
| TMP | 1.028 | 0.163 | 0.873 | 14 | 0.662 | 1.393 |
| NTR | 1.122 | 1.010 | 0.330 | 14 | 0.863 | 1.380 |
| GEN | 1.557 | 2.300 | 0.037 | 14 | 1.038 | 2.077 |
| ERY | 1.563 | 2.999 | 0.010 | 14 | 1.160 | 1.965 |
| TOB | 1.636 | 1.717 | 0.108 | 14 | 0.842 | 2.430 |
| **STR** | **0.264** | **-12.758** | **9.99E-09** | **13** | **0.139** | **0.388** |
| **LVX** | **19.925** | **9.837** | **1.14E-07** | **14** | **15.798** | **24.051** |

**SI Table 3.** The comparison of the mean fold change of IC_50_ values determined at 37°C and at 42°C. Comparisons in **bold** indicate a significant difference between fold changes—meaning the two-fold-change values are statistically significantly different, indicating differential effects of adaptation at different temperatures—after performing a Bonferroni multiple comparison correction (Bonferroni corrected *α* = 0.004). Interestingly, most antibiotic show consistent effects of adaptation across these two temperatures.

| Drug | 37°C Mean Fold Change | 42°C Mean Fold Change | t-statistic | p-value |
| --- | --- | --- | --- | --- |
| AMP | 0.1973 | 0.1366 | 2.062 | 0.0498 |
| **CLI** | **0.4304** | **0.1985** | **3.478** | **0.0019** |
| CPR | 0.4287 | 0.4003 | 0.082 | 0.9358 |
| ERY | 0.9104 | 1.5625 | -3.041 | 0.0061 |
| **FOX** | **0.4188** | **0.7487** | **-3.400** | **0.0022** |
| GEN | 1.3967 | 1.5573 | -0.502 | 0.6198 |
| **LVX** | **6.3127** | **19.924** | **-5.900** | **4.17E-06** |
| NTR | 0.6983 | 1.1216 | -2.786 | 0.0098 |
| STR | 0.2940 | 0.2637 | 0.3263 | 0.7473 |
| TET | 0.2238 | 0.3114 | -2.207 | 0.0367 |
| TMP | 0.5926 | 1.0277 | -2.130 | 0.0436 |
| TOB | 1.0303 | 1.6357 | -1.600 | 0.1303 |

